# ProtoNetStack for DNA-Encoded Source Routing and Majority Aggregation in Protocell Molecular Nanonetworks

**DOI:** 10.64898/2026.08.14.744843

**Authors:** Arman Ferdowsi

## Abstract

Protocell communities can support programmable molecular nanonetworks, yet most demonstrations use broadcast diffusion or fixed sender-receiver circuits. We introduce ProtoNetStack, a network-layer abstraction in which a logical DNA-encoded packet carries a payload, a processing-address list, and an optional forwarding budget. The list determines where localized molecular services transform the packet, not its bidirectional diffusive trajectory.

We formulate a finite-state reaction-transport model whose concentration dynamics and single-copy continuous-time Markov chain use the same generator. Under ideal specificity, positive rates, connected transport, no degradation, and sufficient budget, packet stages advance only in the encoded order and delivery occurs almost surely. All injected concentration is delivered asymptotically. Uniform first-order degradation makes delivery probability the Laplace transform of the lossless delivery-time distribution. A union-bound result separates endpoint delivery from route-faithful delivery under off-target processing.

As an application, we develop cancellation-based strict-majority aggregation on rooted protocell trees. Conservation of token imbalance proves asymptotic correctness and yields a finite-time certificate. With one initial token per node, outside-root mass below one guarantees the correct root sign. Direct matrix-exponential calculations show sequential processing, branching addressability, route-length attenuation, and bounded forwarding work. A 16-condition finite-copy benchmark with 20,000 trajectories per condition shows that off-target reactions can increase endpoint arrival while decreasing route-faithful delivery. Adaptive ordinary differential equation simulations on trees up to 511 compartments show decision time increasing approximately with maximum tree depth and quantify bias from asymmetric loss. ProtoNetStack is therefore a formally analyzable molecular networking architecture and an experimentally testable blueprint. Sequence-resolved gates and chassis calibration remain future work.

## 1. Introduction

Molecular communication provides a way for synthetic or biological microscale agents to exchange information through chemical carriers rather than electromagnetic waves [15, 27]. The networking problem begins once more than one transmitter and one receiver must share this medium. A useful molecular nanonetwork then needs message representation, addressing, forwarding, fault semantics, and interfaces between communication and local computation. These concerns are central to the scope of nano communication engineering, but they are not yet standard design abstractions in synthetic cell communities.

Bottom-up synthetic biology offers unusually direct control over this problem. Protocells and related cell mimics can localize reaction networks, exchange short DNA signals, and be arranged into designed spatial structures [11, 17, 23]. DNA strand displacement provides programmable signal conversion inside such compartments [26, 35]. Experiments have demonstrated DNA communication among proteinosomes, light-controlled sender-receiver systems, amplified intercellular signals, adaptive prototissues, member-specific interactions, and bidirectional interfaces with living cells [3, 14, 17, 20, 25, 32]. These results establish key physical ingredients for molecular networking.

The remaining limitation is architectural. Many synthetic multicellular circuits still encode communication in a fixed sender-receiver wiring pattern. A new logical interaction often requires a new signal family, a new receiver, or a new spatial layout. Diffusion also remains physically broadcast even when only one compartment should process a message. This fixed coupling between the signal species and the receiver response makes it difficult to reuse one communication substrate across several application protocols [21].

Prior work in nano communication networks has introduced protocol stacks based on DNA and enzyme computing [29], molecular addressing by location [22], molecule-level frame representations [13], network models for bacterial colonies [7], simulation platforms [16], and routing schemes for highly constrained nanonetworks [28]. DNA tile systems have also moved computation into message molecules [19], while recent work has considered communication protocols specifically based on DNA [18]. These studies motivate a clean separation between message format, transport, local processing, and application behavior.

This paper develops that separation for compartmental protocell molecular nanonetworks. We introduce ProtoNetStack, a source-routing abstraction in which a DNA-encoded packet exposes one processing address at a time. A compartment whose local address matches the exposed address performs Pop, the logical operation that removes the exposed address from the route state and advances the packet to its next stage. The design relies on reaction specificity rather than directional transport. A packet may physically diffuse through nonaddressed compartments, but it can be transformed only at the compartment selected by its current logical stage.

This distinction prevents a common but consequential ambiguity. In ProtoNetStack, a route is an ordered sequence of processing locations. It is not a claim that an individual molecule follows a unique geometric path. The architecture therefore remains compatible with bidirectional diffusion, permeation, and other passive exchange mechanisms.

The contributions are as follows.

- We define a molecular network-layer service with logical packet states, localized source-route processing, terminal delivery, and an independent forwarding budget.
- We give an exact finite-state model that unifies deterministic concentration dynamics and finite-copy stochastic routing. We prove route-order safety and asymptotically complete lossless delivery.
- We replace a worst-case transit-time approximation by an exact degradation identity. Uniform first-order loss turns delivery probability into the Laplace transform of the lossless delivery time. We also separate endpoint delivery from route fidelity under off-target processing.
- We design distributed cancellation consensus (DCC), a cancellation-based strict-majority aggregation and dissemination protocol on rooted protocell trees. Its proof uses a conserved global imbalance and yields an explicit finite-time correctness certificate.
- We provide reproducible numerical experiments based on sparse matrix exponentiation, finite-copy continuous-time Markov simulation, and adaptive high-order ordinary differential equation integration. The experiments test route semantics, forwarding-budget behavior, finite-copy nonidealities, majority convergence, scaling, and asymmetric loss.

To conclude this section, it is worth noting that the contribution is a theoretical and computational architecture rather than a completed wet-lab implementation. The service model is intentionally above the sequence level. We identify two possible implementation mappings and state their resource trade-offs explicitly. A compiled DNA transducer library directly realizes a finite set of packet states but scales with the number of route states. A modular detachable-stack design could reuse a smaller family of local Pop gates, but that physical mechanism has not yet been experimentally demonstrated. This boundary is important for interpreting the results. A natural next experiment is a three-compartment line with two localized transducers, terminal delivery, and intermediate route-state reporters.

### 2. Related work and positioning

### 2.1. Molecular nanonetwork architectures and packet abstractions

Early molecular communication work established biochemical carriers as a communication medium and emphasized the need to co-design transmitters, propagation, receivers, and interfaces [15]. Systems-theoretic models subsequently connected biological circuit dynamics to communication behavior [24]. Protocol-stack proposals mapped communication functions to enzyme and DNA computing mechanisms [29]. Earlier network environments also used unique single-stranded DNA (ssDNA) identifiers to support addressed messaging [4]. Other studies introduced molecular addressing based on beacon distances [22], frame identifiers and payloads encoded in molecular structure [13], and computation performed during message assembly [19].

ProtoNetStack is closest to this line of work. Its distinct contribution is a localized reaction-gated source route in a compartmental transport graph, together with a formal route semantics and an application proof. The architecture treats a logical packet state as a network-layer object and states exactly which claims follow from the effective reaction-transport model.

Bacterial molecular communication has also motivated multi-node network abstractions, reliability analysis, and protocol simulation [7, 16]. Routing in electromagnetic nanonetworks addresses different physical constraints, but it highlights the same architectural need to define addresses and forwarding behavior under severe node limitations [28]. Our setting differs because transport remains diffusive and logical direction is created by localized chemical transformations.

### 2.2. DNA communication in synthetic cell communities

Encapsulated DNA circuits have enabled communication and distributed processing in protocell populations [17]. Spatially organized sender-receiver arrays have quantified the effects of permeability, degradation, consumption, and catalytic regeneration on DNA signal range [32]. Additional work has shown programmable reaction-rate control, signal amplification, nested compartment organization, adaptive prototissue formation, and selective communication through light-controlled adhesion [3, 14, 25, 33, 34]. DNA has also supported addressable messaging in living bacterial consortia [21].

These platforms demonstrate communication motifs and local signal processing. They do not yet provide a reusable multi-stage source-routing primitive with a correctness proof. ProtoNetStack abstracts the required service without asserting that one particular protocell chassis already realizes every packet operation.

### 2.3. Majority computation

Majority sensing has been implemented experimentally in synthetic microbial consortia [2]. Exact majority is also a central problem in population protocols and chemical reaction network theory [5]. Recent analyses have studied majority thresholds in competitive population dynamics and in stochastic synthetic consortia [8, 10].

Our application has a different purpose. DCC is a topology-aware aggregation protocol on a fixed rooted tree. It uses lossless upward message service and local annihilation to conserve a global signed token count. The protocol is included to show that the network layer supports a nontrivial distributed application with an explicit end-to-end proof.

## 3. System and service model

### 3.1. Compartmental transport graph

Let *G* = (*V, E*) be a finite undirected graph with *V* = {1, …, *n*}. A vertex represents a well-mixed protocell or cell-mimic compartment. An edge {*i, j*} ∈ *E* means that a packet carrier can be exchanged between compartments *i* and *j*. The effective exchange rate is *D*_*ij*_ = *D*_*ji*_ *>* 0. The graph formed by positive-rate edges is assumed connected unless stated otherwise.

The model is agnostic to whether exchange is caused by permeation through semipermeable shells, a pore, a droplet interface, a microfluidic connection, or another engineered interface. Edge rates summarize the chosen carrier, compartment geometry, and environment. They need not be equal.

### 3.2. Logical route and physical location

A source route is an ordered address sequence

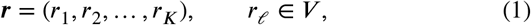

where *r*_*K*_ is the destination. Stages 1 through *K* − 1 require a Pop operation. Stage *K* requires terminal delivery.

A packet state is the pair (*ℓ, i*). The stage *ℓ* identifies the currently exposed logical address *r*_*ℓ*_ The compartment *i* is the current physical location. These quantities have different meanings. Diffusion changes *i* without changing *ℓ*. A correct local processing event changes *ℓ* without requiring a directed physical edge.

#### Remark 1

(Route semantics). The route in Equation (1) specifies an ordered sequence of reaction services. It does not specify the complete physical trajectory of a molecule. A packet can visit a nonaddressed compartment and later leave it unchanged. Route correctness concerns the order and location of packet transformations.

### 3.3. Finite-state reaction-transport process

The state space is

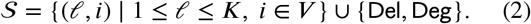

Here De and Deg are absorbing delivered and degraded states. The ideal transition rates are

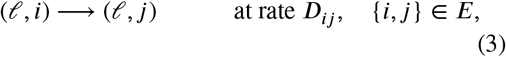

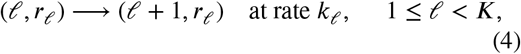

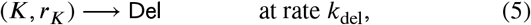

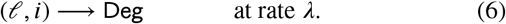

All processing and delivery rates are positive. Ideal specificity means that Equations (4) and (5) are the only transformations that advance or deliver a packet.

Let *Q* be the generator of this finite continuous-time Markov chain (CTMC). A single packet copy has distribution *p*(*t*) satisfying

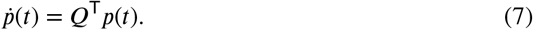

The same equation describes normalized concentrations in a linear reaction-transport model. If an initial concentration *M*_0_ is injected, then *M*_0_*p*_*ℓ,i*_(*t*) is the concentration in state (*ℓ, i*) and *M*_0_*p*_De_ (*t*) is cumulative delivered concentration.

This exact equivalence relies on pseudo-first-order local services. It applies when localized gates are effectively constant over the packet load, when fuel-driven transducers regenerate the active service, or when the equation is used as a calibrated network-level approximation. Finite gate depletion is not represented. A sequence-resolved mass-action implementation must therefore be checked against this service approximation. The main notation is summarized in Table 1.

**Table 1.**
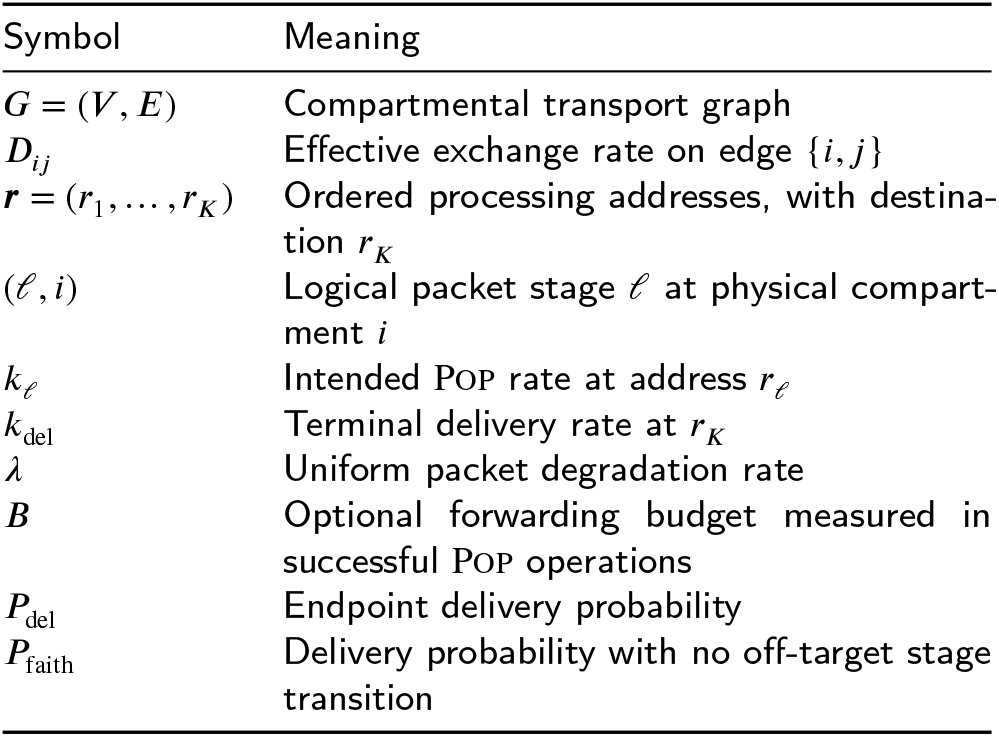
Main notation.

### 4. The ProtoNetStack architecture

### 4.1. Layering

Figure 1 shows the three layers. The physical layer exchanges DNA-compatible carriers over the transport graph. The network layer maintains a logical packet state and exposes localized Pop, delivery, and forwarding-budget services. The application layer interprets the delivered payload. This paper uses majority aggregation as one application, but the network service is independent of that task.

**Figure 1.**
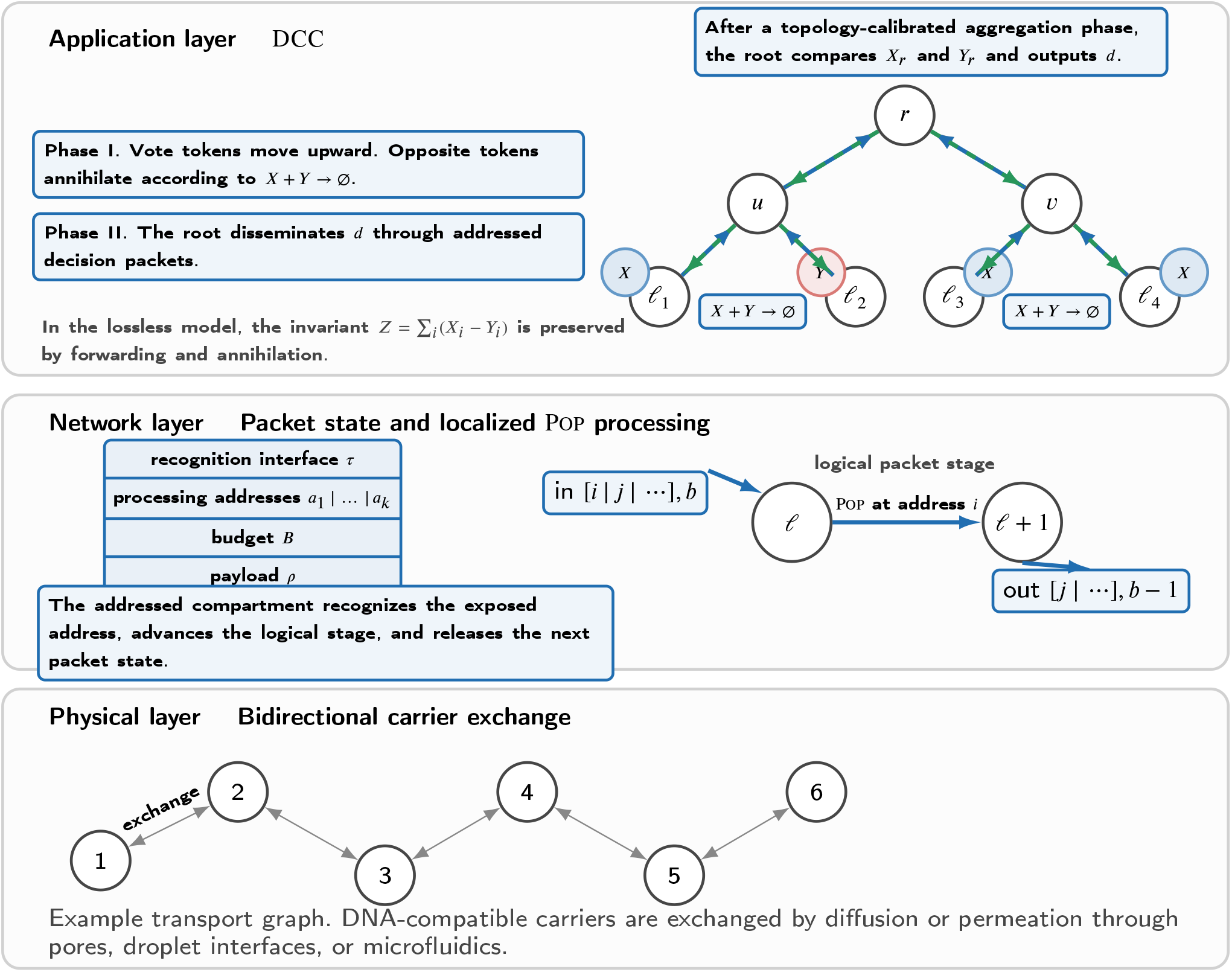
Layered view of ProtoNetStack and its distributed cancellation consensus (DCC) application. The physical layer provides bidirectional carrier exchange. The network layer exposes an ordered processing-address list and a forwarding budget that is consumed only by successful Pop operations. In the lossless reduced model, DCC aggregates a strict majority on a rooted tree and then disseminates the root decision.

### 4.2. Logical packet format

A logical packet is written as

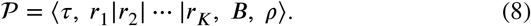

The field *τ* is a recognition interface such as an exposed toe-hold. The ordered addresses define the stages in Equation (1). The optional integer *B* ≥ 0 is a forwarding budget. The payload *ρ* is delivered to local application logic.

Equation (8) is an information-level description. It does not require a monolithic single strand whose prefix is covalently removed. That interpretation would be too restrictive for enzyme-free strand displacement. The model only requires a physical encoding that maps each valid input state to the next logical state and releases a transportable output.

### 4.3. Localized Pop and delivery services

At a nonterminal stage, the addressed compartment performs the effective transformation

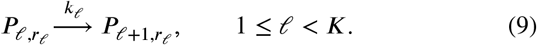

At the destination, the terminal service performs

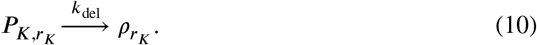

The payload species 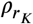 is internal and does not continue as the same network packet.

Two implementation mappings are relevant.

#### Compiled packet-state transducers

A finite route library can assign a distinct DNA signal to each logical suffix. A localized strand-displacement transducer consumes the input signal and releases the next signal. This directly implements Equation (9), but the number of signal and gate designs scales with the number of compiled route states.

#### Modular detachable stack

A packet can instead be represented by a multi-strand assembly whose currently exposed address module is removable. A matching local gate sequesters that module and exposes the next module. Such a design could reuse one address-specific gate family across many suffixes. It would also make carrier size depend on route length and would require experimentally validated connector, permeability, leakage, and release mechanisms. We treat it as a design blueprint rather than an established component.

These mappings implement the same network-layer state transition but have different biochemical resource costs. The formal results do not remove the need for sequence design, secondary-structure screening, crosstalk characterization, and kinetic calibration [26, 31].

### 4.4. Forwarding budget

The optional budget *B* is decremented after each successful Pop operation. In the budgeted model, a transient state is (*ℓ, i, b*), where *b* is the remaining budget, and the intended stage transition is

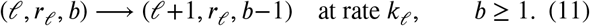

Diffusion and degradation leave *b* unchanged. If the budget reaches zero before stage *K*, the packet becomes nonprocessible. It may continue to diffuse and degrade, but it cannot advance. Terminal delivery does not consume a budget unit. Because every successful ideal stage advance consumes one unit, *b* = *B* − *ℓ* + 1 before stage *ℓ* for a packet initialized at stage one. The budget coordinate can therefore be eliminated when only ideal forwarding is simulated, as in the reduced generator used in the numerical experiments.

The budget is deliberately independent of the address stack. It limits the number of processing reactions and bounds chemical work for malformed or unexpectedly long routes. A fixed finite route already contains only finitely many stages, so the budget is not needed to prove termination. It also does not stop physical diffusion. Calling it a general loop-prevention mechanism would therefore be inaccurate.

## 5. Formal properties of the routing layer

### 5.1. Lossless route execution

#### Theorem 1

(Route-order safety and complete lossless delivery). *Assume that G is finite and connected, all edge rates on E are positive, all intended processing and delivery rates are positive, ideal specificity holds*, λ = 0, *and B* ≥ *K* − 1 *when a forwarding budget is used. Then the following statements hold for every injected packet*.

i. *The stage index never decreases and can change from ℓ to ℓ* + 1 *only in compartment r*_*ℓ*_.
ii. *Delivery can occur only from stage K in compartment r*_*K*_.
iii. *The packet reaches* De *with probability one in finite time*.
iv. *In the deterministic concentration model, the cumulative delivered mass converges to the total injected mass as t* → ∞

*Proof*. Statements (i) and (ii) follow directly from the transition set in Equations (3) to (5). Diffusion changes only physical location. The only stage-changing transition from stage *ℓ< K* is the intended Pop event at *r*_*ℓ*_, and the only delivery transition is at (*K, r*_*K*_).

Fix a stage*ℓ* and add an absorbing success state to the finite location chain. From every physical compartment there is a positive-rate path to *r*_*ℓ*_, followed by a positive-rate processing transition. Finiteness of the state space and positivity of all rates on these paths imply that there are constants *τ* > 0 and *η* > 0 such that, from every location, the probability of completing the stage within time *τ* is at least *1*. By the Markov property, the probability that the stage remains incomplete after *mτ* is at most (1 − *η*)^*m*^. The stage-completion time is therefore almost surely finite and has finite mean. Induction over the finite number of stages proves (iii).

Equation (7) is the Kolmogorov forward equation of the same chain. The delivered concentration divided by the injected concentration equals the absorption probability by time *t*. Statement (iii) implies that this probability converges to one, which proves (iv).

#### Corollary 1

(Forwarding budget). *Under the assumptions of Theorem 1, a budget B* ≥ *K* − 1 *does not change delivery correctness. If B < K* − 1, *terminal delivery is impossible in the ideal model. In the budgeted model, the number of successful stage-advance operations per packet is at most B*.

### 5.2. Uniform degradation

A deterministic upper bound on the completion time is generally unavailable under diffusive transport and exponential reaction waiting times. The correct attenuation statement is distributional.

#### Proposition 1

(Exact attenuation identity). *Let T be the lossless delivery time under the assumptions of Theorem 1. Add a uniform first-order degradation rate λ*≥ 0 *in every transient packet state. Then*

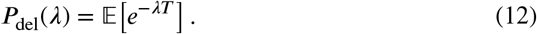

*If E* [*T*] *<* ∞, *then*

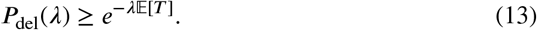

*Proof*. For *λ* = 0, Equation (12) is immediate. For *λ >* 0, couple the lossless routing process to an independent degradation clock *D* ~ Exp(*λ*). Delivery succeeds exactly when *T < D*. Conditioning on *T* gives ℙ(*D > T* I *T*) = *e*^− *λT*^. Taking expectations proves Equation (12). The function *t* ↦ *e*^−*λt*^ is convex, so Jensen’s inequality gives Equation (13).

The identity shows that route attenuation depends on the full delivery-time distribution. Mean delay alone yields only the conservative lower bound in Equation (13). Carrier length, protection, topology, and gate design enter through their effects on *T* and *λ*.

### 5.3. Off-target processing and route fidelity

Endpoint arrival is not sufficient to establish correct route execution. An off-target reaction can advance the logical stage at the wrong compartment and still allow later delivery. Let *D*_*t*_ denote endpoint delivery by time *t*, and let *F* denote at least one incorrect stage transition before absorption. We define

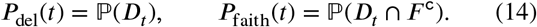

The following bound assumes that terminal delivery itself remains address specific.

#### Proposition 2

(Compositional route-fidelity bound). *Let F*_*ℓ*_*be the event that the ℓth* Pop *operation is processed incorrectly. If* ℙ(*F*_*ℓ*_) ≤*p*_*ℓ*_ *for* 1 ≤*ℓ < K, then for every time horizon*

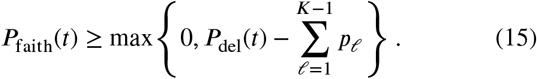

*No independence assumption is required*.

*Proof*. Since 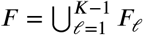, the union bound gives ℙ (*F*) ≤ ∑ _*ℓ*_ *p*_*ℓ*_. Therefore ℙ *D*_*t*_ ∩ *F*^c^) = ℙ (*D*_*t*_) − ℙ (*D*_*t*_ n *F*) ≥ *p*_del_(*t*)− ℙ (*F*). The result follows after truncation at zero.

## 6. Cancellation-based majority aggregation

### 6.1. Task and reduced edge-service model

Let *T* = (*V, E*_*T*_) be a finite rooted tree with root *r*. Each node *i* has an input bit *b*_*i*_ ∈ {0, 1}. We assume a strict majority, so ∑_*i*_ *b*_*i*_ ≠ *n*/2. The goal is for the root to identify the majority and for every node to receive the same decision.

A bit-one vote is represented by token *X* and a bit-zero vote by token *Y*. In node *i*, opposite tokens annihilate according to

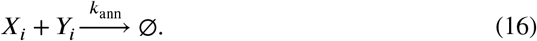

Each nonroot node forwards tokens to its parent *π*(*i*) through the effective services

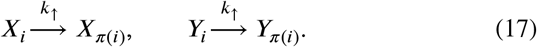

Equation (17) is a reduced application-level service. A ProtoNetStack realization emits a one-edge addressed packet whose payload is decoded into the corresponding token at the parent. The reduced model assumes that this message service is lossless and summarizes its latency by *k*_↑_. It is not a simultaneous packet-level simulation of every in-flight molecule.

The concentration dynamics are

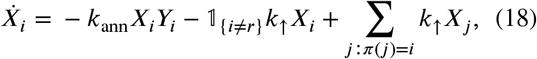

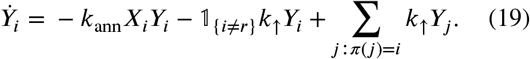

Initialization places one token at each node.

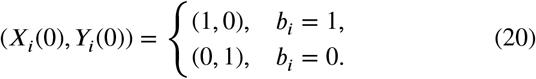

#### Algorithm 1

Cancellation-based strict-majority aggregation and dissemination

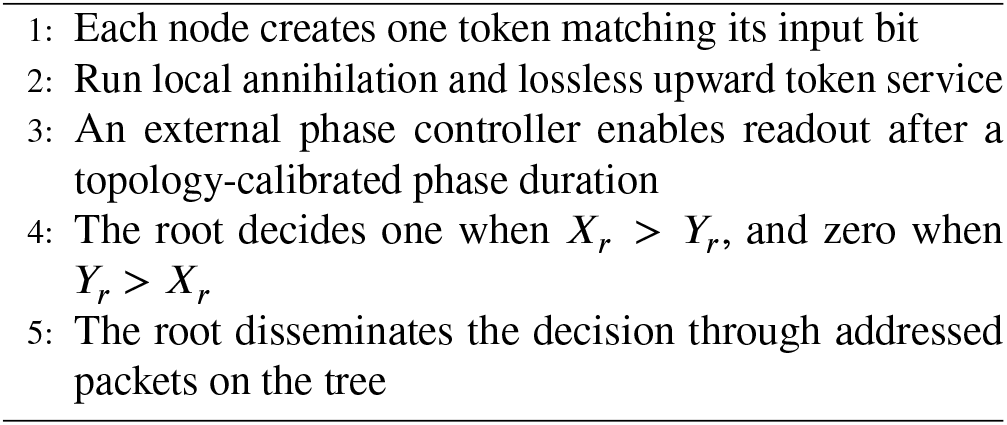

The phase boundary in Algorithm 1 is external. The global outside-root mass used below is an analysis observable, not a quantity that the root can directly measure without additional completion signaling. An implementation can use the topology-dependent bound derived below, external optical gating, or acknowledgements. Autonomous termination detection is outside the present protocol.

### 6.2. Invariant and convergence

Define the global token imbalance and outside-root mass by

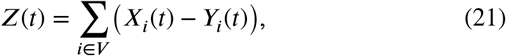

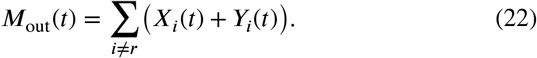

Under one-token initialization, *Z*(0) equals the number of one votes minus the number of zero votes.

#### Theorem 2

(Correct strict-majority aggregation). *Assume k*_ann_ *>* 0, *k*_*↑*_ *>* 0, *and no token loss. Then*

i. *Z*(*t*) = *Z*(0) *for all t* ≥ 0.
ii. *M*_out_(*t*) → 0 *as t* → ∞.
iii. *If Z*(0) *>* 0, *then* (*X*_*r*_(*t*), *Y*_*r*_(*t*)) → (*Z*(0), 0). *If Z*(0) *<* 0, *then* (*X*_*r*_(*t*), *Y*_*r*_(*t*)) → (0, −*Z*(0)).
iv. *At every time satisfying M*_out_(*t*) *< Z*(0), *the sign of X*_*r*_(*t*) − *Y*_*r*_(*t*) *equals the sign of Z*(0).

*Proof*. An annihilation event removes one *X* and one *Y*, so it preserves their difference. Upward service removes a token from one node and adds the same token to its parent. Summing over all nodes cancels every forwarding term. This proves (i).

For a leaf *i*, its total mass *M*_*i*_ = *X*_*i*_ + *Y*_*i*_ satisfies 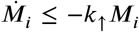, hence *M*_*i*_(*t*) → 0. Consider a nonroot node whose children have already been shown to converge to zero. Its total mass obeys

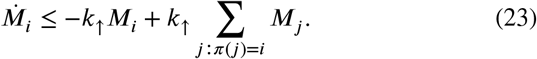

The input term converges to zero. A comparison argument implies *M*_*i*_(*t*) → 0. Induction from the leaves toward the root proves (ii).

Let *z*_*r*_ = *X*_*r*_ − *Y*_*r*_. By (i),

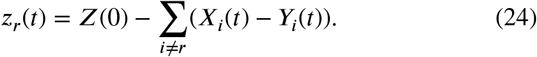

Statement (ii) implies *z*_*r*_(*t*) → *Z*(0). If *Z*(0) *>* 0, then *X*_*r*_ is bounded away from zero for sufficiently large time. The root equation for *Y*_*r*_ consists of the negative term −*k*_ann_*X*_*r*_*Y*_*r*_ and an input from children that converges to zero. Standard comparison with a stable linear equation gives *Y*_*r*_(*t*) → 0, and then *X*_*r*_(*t*) → *Z*(0). The negative case is symmetric. This proves (iii).

Finally,

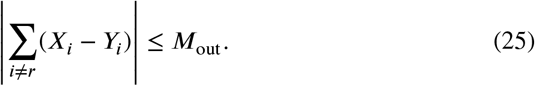

If *Z*(0) *>* 0 and *M*_out_ *< Z*(0), Equation (24) gives *z*_*r*_ *>* 0. The case *Z*(0) *<* 0 is symmetric. This proves (iv).

#### Corollary 2

(Universal finite-time threshold). *With one token per node and a strict majority, |Z*(0) > 1|. *Therefore any readout time with M*_out_(*t*) *<* 1 *guarantees the correct root decision in the lossless model. A threshold E <* 1 *is valid without knowing the initial majority margin*.

#### Proposition 3

(Topology-calibrated phase duration). *Let d*_*i*_ *be the number of tree edges from node i to the root and let H* = max_*i*_ *d*_*i*_. *Under the assumptions of Theorem 2 and the one-token initialization*,

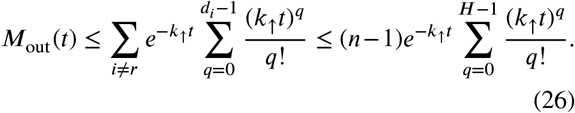

*Consequently, any externally chosen phase duration for which the rightmost expression is below one guarantees a correct strict-majority readout for every input assignment*.

*Proof*. Let *M*_*i*_ = *X*_*i*_ + *Y*_*i*_. Removing annihilation from the dynamics deletes the nonpositive term −2*k*_ann_*X*_*i*_*Y*_*i*_. Comparison for the resulting positive linear system therefore upper-bounds each *M*_*i*_(*t*) by the mass in a pure upward-transport process. A unit initially at depth *d*_*i*_ remains outside the root at time *t* exactly when a sum of *d*_*i*_ independent exponential waiting times of rate *k*_↑_ exceeds *t*. Its survival probability is the Erlang tail shown in the first sum of Equation (26). Summing over initial nodes proves the first inequality. The Erlang tail is nondecreasing in the integer shape parameter, which gives the second inequality. The final statement follows from Corollary 2.

#### Corollary 3

(Agreement after dissemination). *If the root decision is correct and every addressed decision packet on the finite tree is eventually delivered without corruption, then every node eventually adopts the same correct decision*.

*Proof*. The root sends to its children. Every node that first accepts the decision sends it to its own children. Induction on tree depth completes the proof.

### 6.3. Symmetric and asymmetric token loss

Add first-order token degradation rates *µ*_*X*_ and *µ*_*Y*_. Forwarding and annihilation still cancel in the derivative of the global imbalance, giving

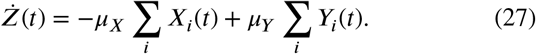

If *µ*_*X*_ = *µ*_*Y*_ = *µ*, then Ż = −*µZ* and

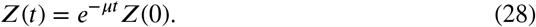

The sign is preserved at every finite time. This mathematical fact does not guarantee a robust experimental readout because the absolute signal and the margin above detection noise also decay. When *µ*_*X*_ ≠ *µ*_*Y*_, Equation (27) contains a composition-dependent bias and no sign invariant remains.

## 7. Numerical methods

This section specifies the numerical solvers, benchmark designs, parameter values, and uncertainty calculations used in the computational study. The default parameter values are summarized in Table 2. All experiments are dimensionless design studies^1^. They are not fitted to a particular protocell chassis.

**Table 2.** Default dimensionless numerical parameters.

| Parameter | Meaning | Value |
| --- | --- | --- |
| $D$ | Default edge exchange rate | 0.2 |
| $k_\ell$ | Intended POP rate | 1 |
| $k_{\text{del}}$ | Delivery rate | 1 |
| $\lambda$ | Packet degradation rate | 0.05 |
| $k_{\text{ann}}$ | Token annihilation rate | 10 |
| $k_\uparrow$ | Upward token service rate | 1 |
| $\epsilon$ | Outside-root decision threshold | 0.05 |

### 7.1. Routing calculations

The linear routing systems were solved by applying a sparse matrix exponential to the generator in Equation (7) [1]. This avoids time-discretization error and concentration clipping. The default rates were *D* = 0.2, *k*_*ℓ*_ = 1, *k*_del_ = 1, and *λ* = 0.05.

The line experiment used eight compartments and the route (2, 3, 4, 5, 6, 7, 8). The route-length experiment used line networks requiring one through ten Pop operations. For each length, we computed the exact eventual delivery probability under degradation, the mean lossless delivery time, and the Jensen lower bound from Equation (13).

The branching graph contained edges 1 ↔ 2 ↔ 3 ↔ 5 and 2 ↔ 4 ↔ 6. Two logical packets used routes (2, 3, 5) and (2, 4, 6). Edge rates on the first branch were 0.22, while rates on the second branch were 0.16.

The forwarding-budget experiment used a four-node ring and the repeated-address route (2, 3, 4, 1, 2). We compared a sufficient budget of four with an insufficient budget of two. Cumulative Pop work was obtained by integrating the intended transition flux.

### 7.2. Finite-copy stress test

A single-packet CTMC was simulated exactly using the Gillespie direct method [12]. Although efficient surrogate trace generation has been studied for transcription models [9], all trajectories here are exact. Each trajectory independently drew edge, Pop, and delivery rates from lognormal distributions with the stated nominal means and coefficient of variation (CV) in {0, 0.1, 0.2, 0.3}. At a nonaddressed compartment, an off-target stage transition occurred at rate *ϵk*_*ℓ*_, with *ϵ* ∈ {0, 0.01, 0.02, 0.05}. The degradation rate remained 0.05.

The full factorial design used 20,000 trajectories for each of the 16 conditions, route (2, 3, 5), and horizon *t* = We recorded endpoint delivery, route-faithful delivery, any off-target event, and conditional median delivery time. Binomial intervals are 95 percent Wilson intervals [30]. The deterministic nominal value at *t* = 40 was computed from the same generator.

### 7.3. Majority aggregation calculations

The nonlinear system in Equations (18) and (19) was integrated with the adaptive DOP853 explicit Runge-Kutta method of order eight. The illustrative and scaling runs used relative tolerances between 5 × 10^−11^ and 2 × 10^−8^ and absolute tolerances between 10^−13^ and 10^−10^. The batched asymmetric-loss study used relative tolerance 10^−7^ and absolute tolerance 10^−9^. No nonnegativity clipping was applied. Default rates were *k*_ann_ = 10 and *k*_↑_ = 1.

The operational decision time was the first time at which *M*_out_ ≤ 0.05. By Corollary 2, this threshold certifies correctness for every strict-majority one-token instance in the no-loss model. It remains an analysis observable rather than a local stopping rule.

Scaling was evaluated on full binary trees with *n* ∈ {15, 31, 63, 127, 255, 511}. Each size used 24 independently sampled minimal-margin assignments with *Z*(0) = 1 and a root input of zero. We report medians and interquartile ranges.

For the loss experiment, *n* = 63 and *µ* = 0 05. We set *µ*_*X*_ = (1 + *δ*)*µ* and *µ*_*Y*_ = (1 − *δ*)*µ*, with *δ* ∈ {−0.6, −0.4, −0.2, 0, 0.2, 0.4, 0.6}. We sampled 160 independent Bernoulli assignments and included each complement, giving 320 decisions per value of *δ*. The absolute initial majority margins ranged from one to 21, with median seven. Complement pairing balances the two decision labels. The threshold is an operational readout rule in this asymmetric-loss experiment, not a consequence of the lossless certificate. Confidence intervals were obtained by 4,000 bootstrap resamples at the pair level, preserving within-pair dependence.

## 8. Results

### 8.1. Source-route processing is sequential even though transport is diffusive

Figure 2 shows the stage-local concentration at each addressed node on the eight-compartment line. Each trace is normalized by its own maximum so that ordering remains visible despite degradation. Peak times increase from 1.55 at node 2 to 36.2 at node 8. Peak amplitudes in the unnormalized data decrease from 9.83 × 10^−2^ to 3.48 × 10^−4^. The result matches the route semantics in Theorem 1. Physical diffusion is not directed, but stage transformations occur in the prescribed order.

**Figure 2.**
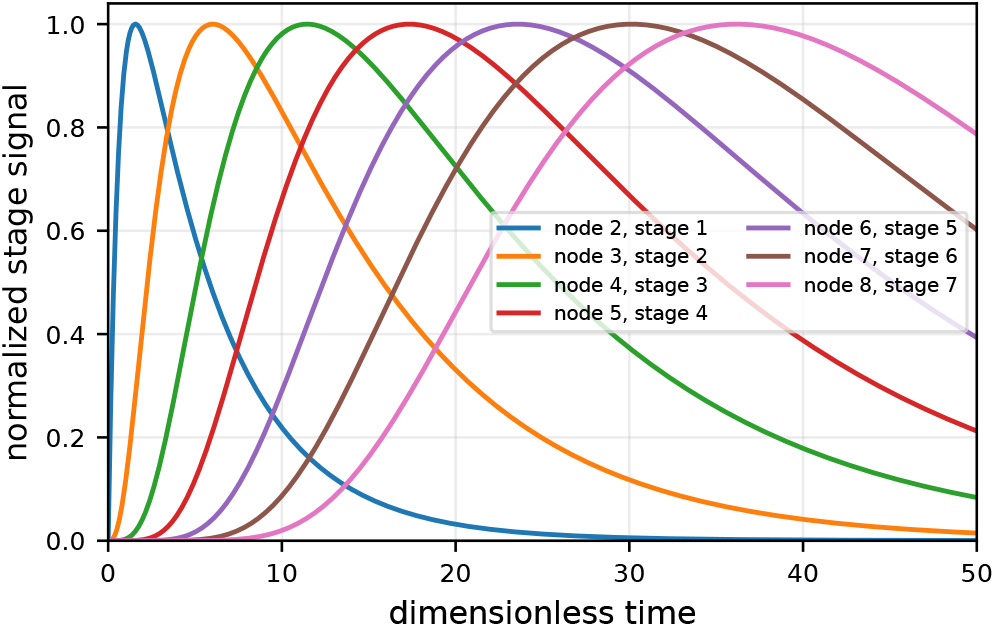
Sequential logical processing on an eight-compartment line. Each stage-local trace is divided by its own maximum. The normalization reveals the ordered peak times, while the raw data retain the route-length attenuation.

The route-length experiment in Figure 3 quantifies the cost of repeated diffusive search and processing. In this line geometry, the mean lossless delivery time rises from 21 for one Pop operation to 462 for ten. The exact delivery probability at *λ* = 0.05 falls from 0.439 to 1.05 × 10^−3^. The Jensen expression remains a valid lower bound but becomes highly conservative for long routes. This gap confirms that the complete delivery-time distribution matters, as stated in Proposition 1.

**Figure 3.**
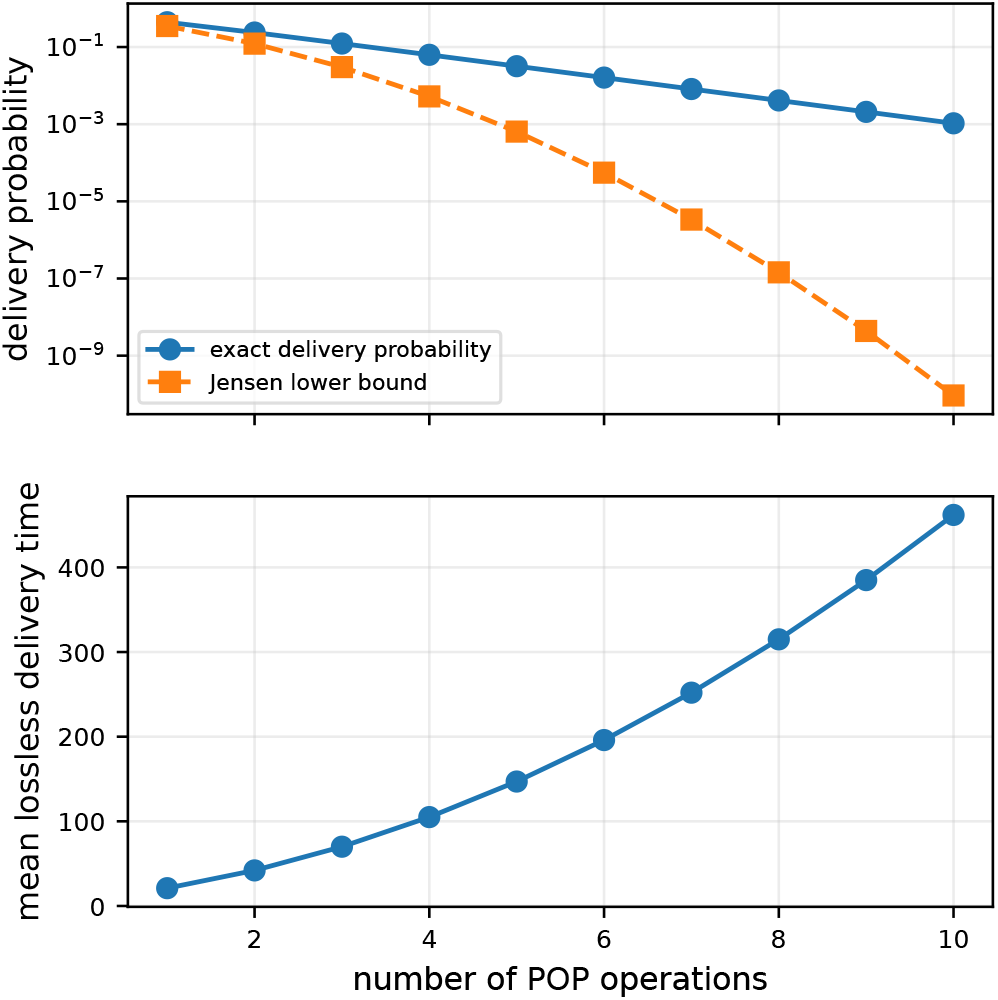
Route-length effects on a line. The upper panel compares exact eventual delivery under degradation with the Jensen lower bound. The lower panel shows mean delivery time in the corresponding lossless process.

### 8.2. One physical graph supports distinct logical destinations

Two packets injected at node 1 share their first address and then diverge toward nodes 5 and 6. Figure 4 shows cumulative delivery at the intended endpoints. At *t* = 45, the delivered fractions are 0.170 and 0.130. The difference is caused by the lower exchange rates on the second branch. The packets use the same physical graph and the same stage semantics, while their headers select different localized transformations.

**Figure 4.**
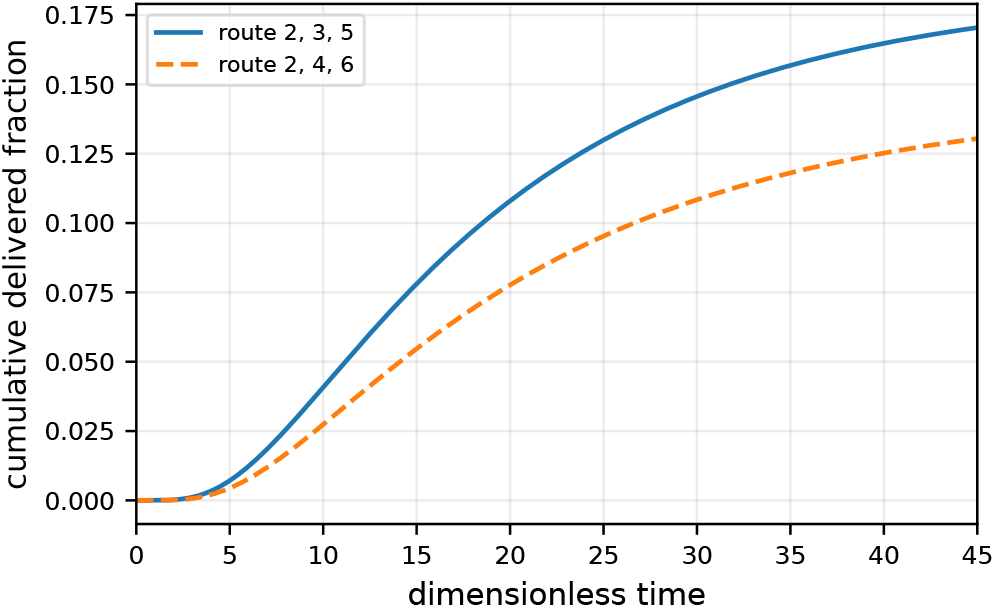
Addressable delivery on a branching transport graph. Two logical routes coexist on the same bidirectional physical substrate. Different edge rates produce different latency and attenuation.

This result demonstrates logical addressability, not geometric confinement. A route-*A* molecule can diffuse onto the other branch before returning. It remains route faithful because only nodes 2, 3, and 5 can transform or deliver its logical states.

### 8.3. The forwarding budget limits processing work

Figure 5 uses a route that revisits previously addressed nodes. A budget of four permits all required Pop operations. A budget of two prevents terminal delivery and moves probability mass into an expired nonprocessible stage. At *t* = 50, the insufficient-budget expired-state mass is 0.076 after additional mass has degraded.

**Figure 5.**
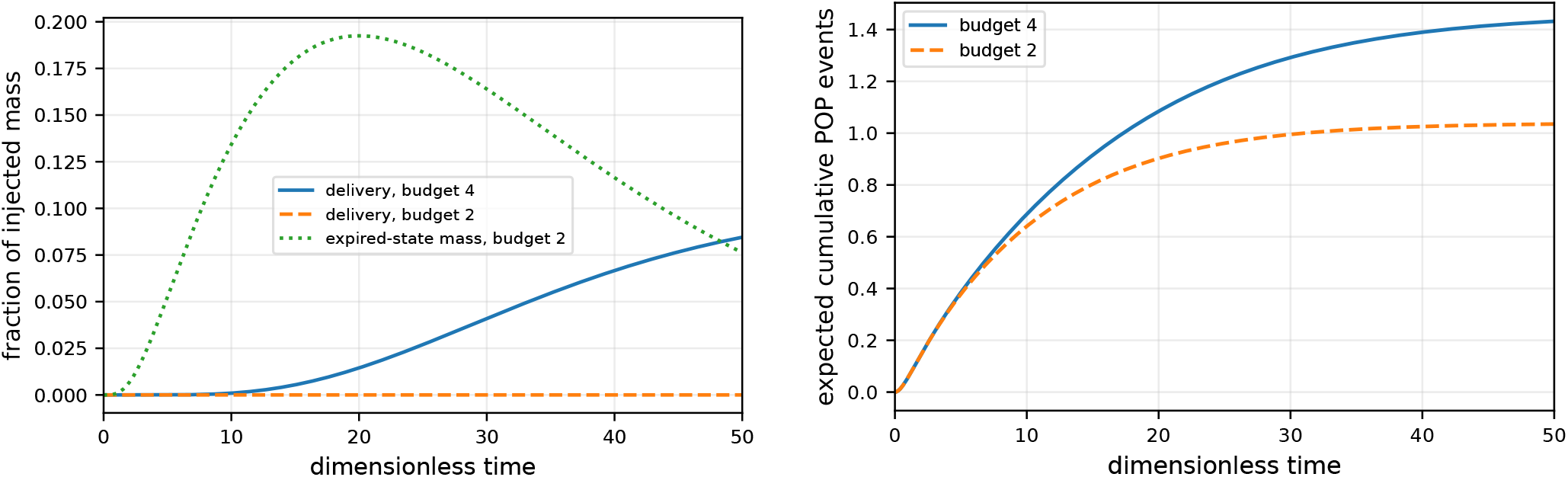
Forwarding-budget behavior on a cyclic transport graph. The left panel shows delivery and expired-state mass. The right panel shows expected cumulative intended Pop events. A small budget truncates logical processing but does not stop physical diffusion.

By *t* = 50, the expected cumulative numbers of Pop events are 1.43 for budget four and 1.04 for budget two under the chosen degradation regime. These values are below the nominal budgets because many packets degrade before using every available operation. The experiment supports the intended interpretation. The budget bounds transformation work, while expired packets can still diffuse.

### 8.4. Endpoint arrival can conceal route errors

The nominal finite-copy result agrees with the deterministic model. At *t* = 40, the matrix-exponential prediction is 0.1648. The CTMC estimate from 20,000 trajectories is 0.1633, with 95 percent Wilson interval [0.1582, 0.1685].

Figure 6 separates endpoint delivery from route-faithful delivery. Increasing the off-target rate can raise raw endpoint arrival because a spurious stage advance acts as a shortcut. At zero kinetic heterogeneity, raising *ε* from zero to 0.05 increases endpoint delivery from 0.1633 to 0.1931, while route-faithful delivery falls to 0.1012. At coefficient of variation 0.3, the corresponding endpoint and route-faithful probabilities are 0.1789 and 0.0930. The probability of at least one off-target event is about 0.38 in all *ε* = 0.05 conditions.

**Figure 6.**
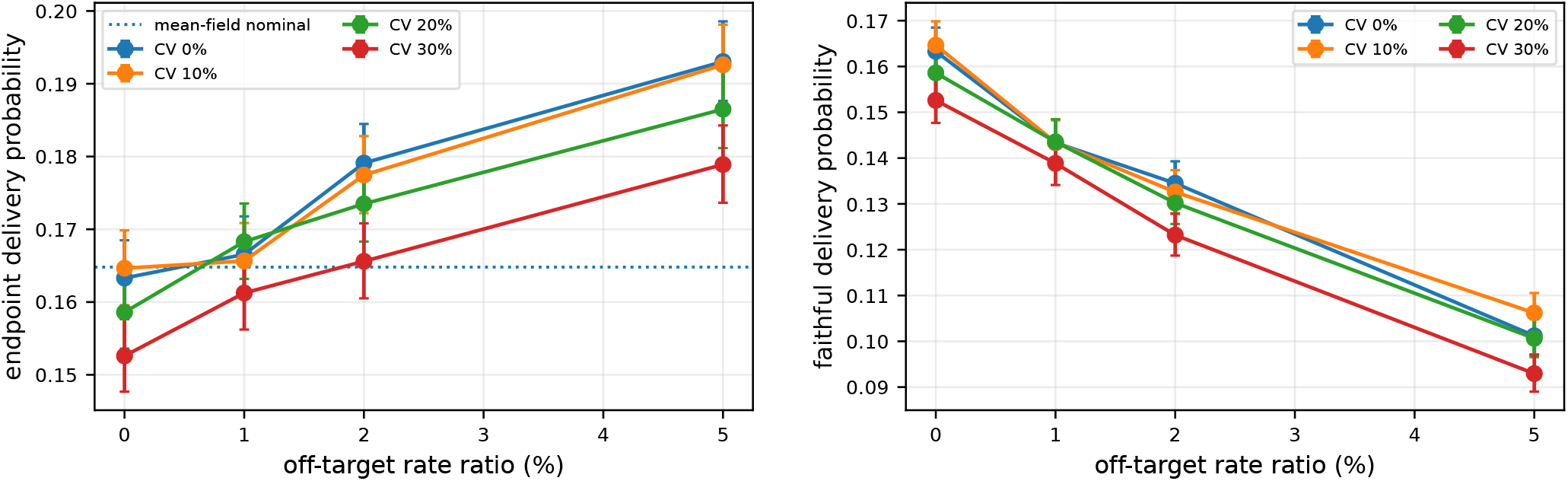
Finite-copy full factorial stress test. The left panel reports endpoint delivery by time 40. The right panel reports route-faithful delivery. Error bars are 95 percent Wilson intervals over 20,000 trajectories per condition. The dotted line on the left is the deterministic nominal prediction.

Kinetic heterogeneity alone has a smaller effect than off-target processing in this experiment. The main semantic failure is therefore not missing the destination. It is reaching the destination after an incorrect sequence of transformations. This is why route-fidelity metrics and the bound in Proposition 2 are necessary.

### 8.5. Cancellation aggregation satisfies the finite-time certificate

Figure 7 shows a 31-node minimal-margin instance with *Z*(0) = −1. Tokens initially spread across the tree, opposite tokens annihilate, and the remaining *Y* mass concentrates at the root. The outside-root mass crosses 0.05 at approximately *t* = 10.78. At that time, the root has *X*_*r*_ ≈ 0.0024 and *Y*_*r*_ ≈ 1.0193, so its sign is already correct. The global imbalance remains −1 within numerical precision.

**Figure 7.**
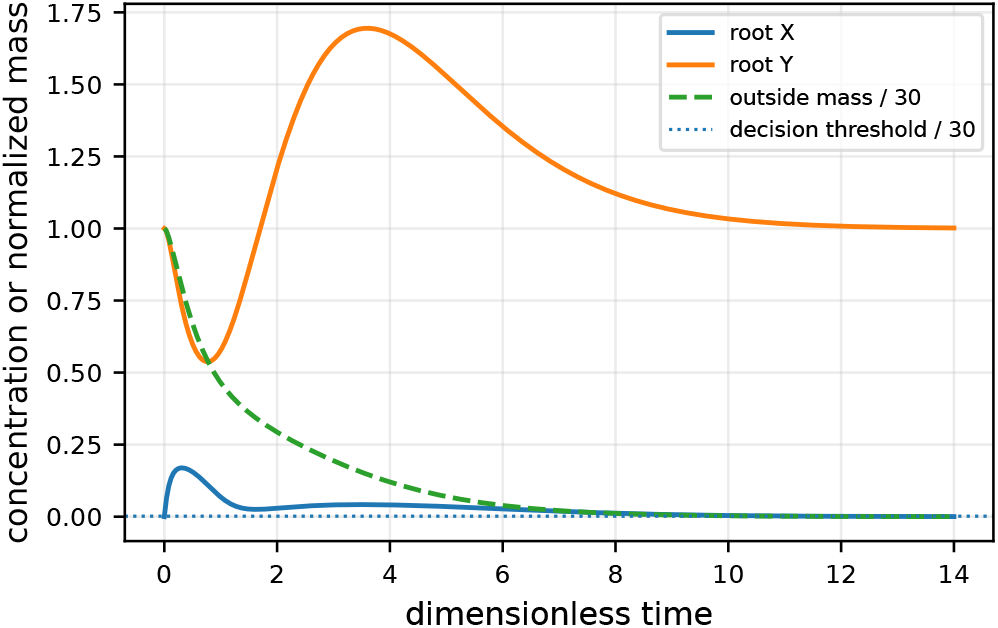
Cancellation-based aggregation on a 31-node full binary tree with strict majority zero. The outside-root mass decays below the decision threshold while the surviving majority token concentrates at the root.

A separate 63-node invariant run with *Z*(0) = 9 has maximum absolute drift 1.07 × 10^−14^. This is solver error rather than a model effect and confirms that adaptive integration preserves the analytical invariant to floating-point precision.

### 8.6. Decision time grows approximately with tree height

Across 24 minimal-margin instances per size, median decision time grows from 8.32 at *n* = 15 to 20.04 at *n* = 511 in the left panel of Figure 8. A linear fit against maximum root-to-leaf depth *H* = log_2_(*n* + 1) − 1 has slope 2.34 and coefficient of determination 0.9999 over the six tested sizes. This is an empirical property of the chosen balanced-tree geometry and rate regime, not a general complexity theorem. It is consistent with sequential transport across increasing tree depth.

**Figure 8.**
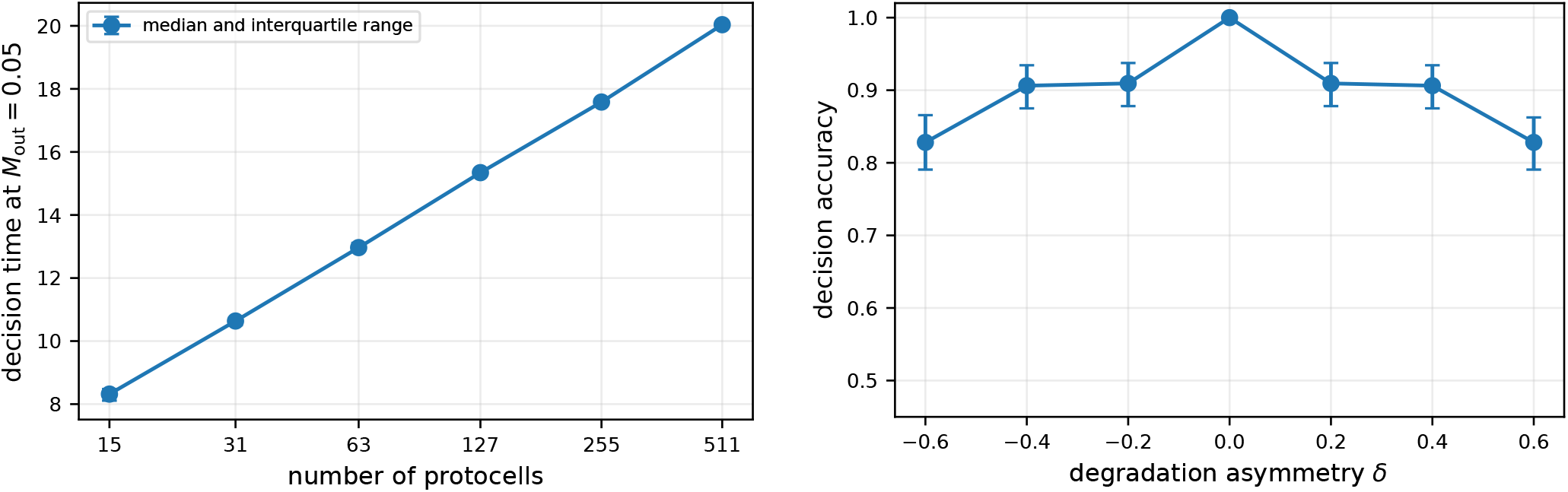
Consensus timing and loss sensitivity. The left panel shows decision time on full binary trees. Points are medians over 24 independently sampled minimal-margin instances and error bars show interquartile ranges. The right panel shows majority accuracy under asymmetric token degradation on 63-node trees. Each point uses 160 assignments and their complements. Error bars are 95 percent pair-bootstrap intervals.

### 8.7. Symmetric loss preserves sign, while asymmetric loss biases decisions

At *δ* = 0, all 320 complement-paired decisions are correct, consistent with Equation (28). Accuracy decreases to 0 909 near |*δ*| = 0 2, 0 906 at |*δ*| = 0 4, and 0 828 at |*δ*| = 0.6. Pair-bootstrap intervals are shown in the right panel of Figure 8. The curve is symmetric because every sampled assignment is paired with its complement.

These results do not contradict sign preservation under symmetric loss. They show that differential degradation changes the signed global quantity itself. They also reinforce the practical limitation of symmetric loss. Even when the sign is mathematically preserved, a real detector must resolve a progressively smaller absolute signal.

## 9. Discussion and experimental roadmap

### 9.1. What the formal model establishes

The routing theorem establishes a precise network-layer guarantee. Under ideal localized services, connected transport, positive rates, no degradation, and sufficient budget, every packet stage is processed in order and all injected mass is eventually delivered. The result allows arbitrary bidirectional physical diffusion. It therefore captures the core advantage of reaction-gated addressing.

The theorem does not establish that a particular DNA sequence set realizes ideal specificity. It also does not imply useful finite-time delivery under severe degradation. The line experiment shows that route delay and attenuation can become prohibitive even though asymptotic lossless correctness is exact. Stabilized carriers, faster exchange, amplification, replication, or shorter logical routes may be required.

The finite-copy study exposes another design requirement. Endpoint detection cannot certify route correctness when off-target reactions can advance a header. This observation complements chemical-reaction-network analyses of coexistence between communication chemistry and surrounding biochemical processes [6]. A future experiment should therefore include intermediate state reporters or chemically encoded provenance markers, not only a terminal fluorescence signal.

### 9.2. Biochemical implementation trade-offs

A compiled transducer implementation is the most direct route to an initial experiment. For a fixed small set of routes, every suffix can be represented by a separate strand and every intended state transition by a localized transducer. This increases the number of sequence designs but avoids the unproven assumption that one generic gate can detach an arbitrary first address while preserving an unknown suffix.

A modular stack is more attractive for scalability. Its key experimental requirements are a stable transportable assembly, an exposed address module, selective irreversible removal, release of a forwardable remainder, and a budget token that can be consumed independently. Longer stacks may diffuse or permeate more slowly. Connector domains may also increase secondary structure and crosstalk. These effects can be represented by stage-dependent *D*_*ij*_, *k*_*ℓ*_, and *λ*, but they must be measured.

Address-space scaling is similarly physical rather than purely combinatorial. DNA sequence space is large, but the number of simultaneously usable domains is limited by orthogonality, leakage, secondary structure, salt conditions, temperature, and chassis-specific transport. Sequence screening and kinetic calibration are therefore part of network provisioning, not an implementation detail that can be ignored.

### 9.3. Minimal validation sequence

A staged experimental program can test the architecture without first implementing the majority application.

1. Demonstrate one localized input-to-output transducer inside a semipermeable compartment and verify that the output remains transportable.
2. Build a three-compartment line with two processing stages and terminal delivery. Measure intermediate state activation, endpoint delivery, leakage, and degradation.
3. Compare two headers on a branching layout. Confirm that endpoint arrival and route-faithful intermediate reporters agree.
4. Add a finite forwarding budget to a repeated-address header and measure both expired-state accumulation and consumed gate or fuel.
5. Only after these network-layer tests, implement token annihilation and a small externally clocked aggregation tree.

The existing protocell literature provides several relevant components, including semipermeable DNA-communicating proteinosomes, anchored strand-displacement gates, light-controlled phase activation, and tunable signal regeneration [17, 32]. The missing experiment is their composition into a stateful multi-stage forwarding service.

### 9.4. Limitations

The current work has some principal limitations. First, no sequence-resolved Pop design is provided. Second, gate depletion and fuel consumption are absorbed into effective first-order rates. Third, well-mixed compartments exclude intracompartmental gradients and crowding. Fourth, the majority simulations use reduced edge service rather than explicit in-flight packet species. Fifth, the phase boundary for root readout is externally controlled. Sixth, all parameters are dimensionless and are not quantitative predictions for a particular platform.

These limitations immediately define the next modeling steps. A chassis-specific model should couple sequence-level strand-displacement kinetics, carrier-dependent permeability, finite gate inventories, stochastic copy numbers, and spatial diffusion. The network abstraction remains useful because each measured component can be mapped to an effective transition or fault parameter and tested against the stated guarantees.

## 10. Conclusion

We introduced ProtoNetStack, a network-layer architecture for DNA-encoded source routing in protocell molecular nanonetworks. Its central principle is reaction-gated logical direction on top of physically bidirectional transport. A route specifies where packet transformations occur, not the complete geometric path of a molecule.

The finite-state model yields exact route semantics, asymptotically complete lossless delivery, a degradation identity based on the delivery-time Laplace transform, and a compositional route-fidelity bound. The DCC application shows how the communication service can support strict-majority aggregation with a conserved invariant and a finite-time correctness certificate.

The numerical results confirm the predicted stage order and majority invariant, quantify route-length attenuation, show how forwarding budgets bound processing work, and demonstrate why endpoint arrival must be separated from route fidelity. They also show moderate height-dependent aggregation time in balanced trees and the bias introduced by asymmetric token degradation.

The architecture is ready for small-scale experimental falsification, not yet for claims of chassis-independent implementation. The most informative next step is a short line of compartments with intermediate route-state reporters. Such an experiment would directly test the distinction between physical diffusion and logical source-route processing that underlies the entire design.

## Acknowledgment

This work was funded in whole by the Austrian Science Fund (FWF) 10.55776/ESP1705325 (STAAC Project).

## Footnotes

1 The source code is hosted in a private GitHub repository and will be made public upon acceptance of the manuscript.

